# Structural basis for ATG9A recruitment to the ULK1 complex in mitophagy initiation

**DOI:** 10.1101/2022.07.12.499634

**Authors:** Xuefeng Ren, Thanh N. Nguyen, Wai Kit Lam, Cosmo Z. Buffalo, Michael Lazarou, Adam Lee Yokom, James H. Hurley

## Abstract

The assembly of the autophagy initiation machinery nucleates autophagosome biogenesis, including in the PINK1- and Parkin-dependent mitophagy pathway implicated in Parkinson’s disease. The structural interaction between the sole transmembrane autophagy protein, ATG9A, and components of the ULK1 complex is one of the major missing links needed to complete a structural map of autophagy initiation. We determined the 2.4 Å x-ray crystallographic structure of the ternary structure of ATG9A C-terminal tail bound to the ATG13:ATG101 HORMA dimer, which is part of the ULK1 complex. We term the interacting portion of the extreme C-terminal part of the ATG9A tail the “HORMA dimer interacting region” (HDIR). This structure shows that the HDIR binds to the HORMA domain of ATG101 by β-sheet complementation such that the ATG9A tail resides in a deep cleft at the ATG13:ATG101 interface. Disruption of this complex in cells impairs damage induced PINK1/Parkin mitophagy mediated by the cargo receptor NDP52.

## Introduction

Macroautophagy (henceforth autophagy) is a conserved cellular degradative process that maintains cellular homeostasis under a variety of stress conditions including starvation, mitochondrial damage, and microbial invasion. One of the major functions of autophagy is to selectively target and degrade various unneeded or dangerous cellular cargoes. Disfunction of selective autophagy and the consequent accumulation of problematic cargoes contribute to a multitude of human disease (Mizushima and Levine 2020). Parkinson’s Disease (PD) is one of the clearest examples of an autophagy defect linked to human disease. In *PRKN* and *PINK1* PD patients, the failure to clear damaged mitochondria via mitophagy is thought to contribute to disease due to the progressive loss of dopaminergic neurons in the substantia nigra pars compacta (Panicker et al. 2021; Singh and Muqit 2020). Selective autophagy proceeds from the recognition of cargoes, formation of autophagy initiation sites, de novo synthesis of a double lipid bilayer termed the phagophore (or isolation membrane), maturation of the phagophore into a closed autophagosome which sequesters cargo, and finally autophagosome-lysosome fusion leading to cargo degradation (Chang, Jensen, and Hurley 2021; Ktistakis 2021; Maruyama et al. 2021). A set of core autophagy initiation proteins bridge cargo recognition to isolation membrane biogenesis and elongation into the cup shaped phagophore (Chang, Jensen, and Hurley 2021; Maruyama et al. 2021). In mammalian autophagy, the Unc-51 like autophagy activating kinase (ULK1) complex, PI3KC3 Complex I (PI3KC3-C1), ATG12-5-16L1 ATG8 conjugation machinery, ATG2A/B, and ATG9A are all fundamental to autophagosome formation (Hurley and Young 2017). How these complexes interact to trigger initiation has been challenging to dissect due to the presence of overlapping and partially redundant activities and the large size and dynamic character of the protein complexes.

The ULK1 complex is comprised of four proteins: the ULK1 kinase, autophagy-related protein (ATG) 13, ATG101 and, focal adhesion kinase family interacting protein of 200 kDa (FIP200) (Ganley et al. 2009; Hosokawa et al. 2009; Jung et al. 2009; Mercer, Kaliappan, and Dennis 2009). The ULK1 and ULK2 kinases regulate much of the autophagy pathway through phosphorylation events (Egan et al. 2015). The non-kinase subunits of the ULK1 complex have numerous scaffolding and bridging roles, some of which act upstream of ULK kinase activity. FIP200 is recruited by autophagic cargo receptors to mark sites of autophagy initiation and scaffolds the assembly of the ULK1 complex (Ravenhill et al. 2019; Shi, Chang, et al. 2020; Turco et al. 2019; Vargas et al. 2019). Cargo receptors including p62 and NDP52 engage with the C-terminal coiled-coil and Claw domains of FIP200 (Ravenhill et al. 2019; Turco et al. 2019; Vargas et al. 2019). These interactions place the ULK1 complex in an upstream position in many forms of selective autophagy, such that it is thought to be responsible for bridging to the rest of the autophagy initiation machinery (Turco, Fracchiolla, and Martens 2020).

ATG13 translocation to initiation sites is a key early step in both starvation-induced autophagy and mitophagy (Dalle Pezze et al. 2021; Zachari et al. 2019). The ATG13 and ATG101 subunits of the ULK1 complex form a heteromeric dimer through homologous Hop1/Rev7/Mad2 (HORMA) domains (Figure 1A) (Qi et al. 2015; Suzuki et al. 2015). HORMA domains contain a ‘safety belt’ which conformationally regulates binding to protein proteins (Gu, Desai, and Corbett 2022). ATG101 contains a protruding Trp-Phe (WF) finger motif, which is important for autophagy initiation (Kim et al. 2018; Qi et al. 2015; H. Suzuki et al. 2015), but whose precise function is unknown. In addition to a HORMA domain, ATG13 contains a C-terminal intrinsically disordered region of ∼300 residues. This IDR is responsible for interaction with FIP200 and the ULK1 kinase (Hieke et al. 2015; Shi, Yokom, et al. 2020; Yamamoto et al. 2016).

**Figure 1.**
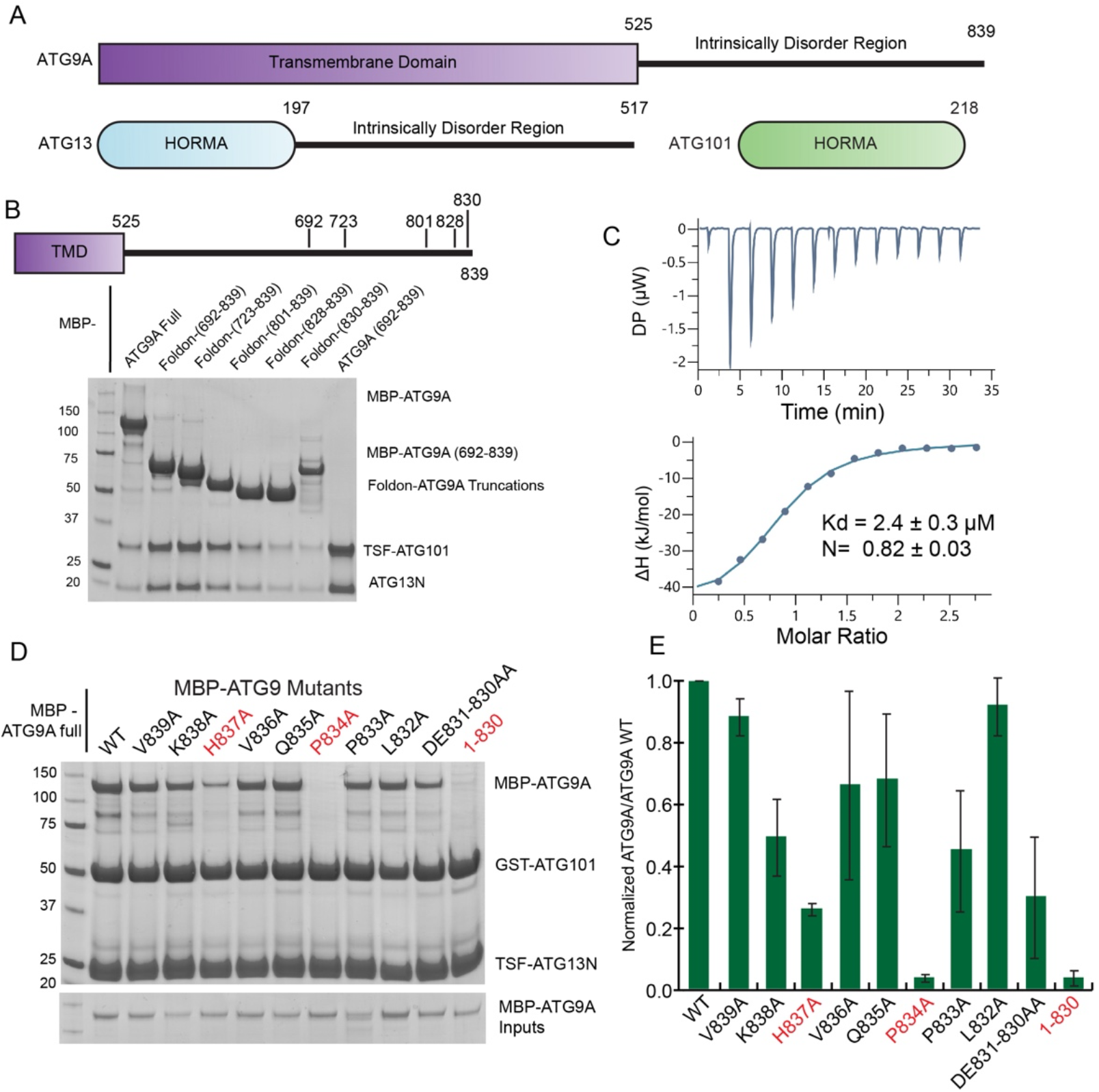
Domain Arrangement and ATG9A C-terminal Tail Pulldowns. A) Domain Schematic of ATG9A, ATG13 and ATG101. B) MBP pulldown assay using recombinant MBP-ATG9A in DDM/CHS micelles, MBP-FOLDON-ATG9A and MBP-ATG9A constructs bound to amylose resin. Incubated with ATG13 ^HORMA^ /TSF-ATG101 and visualized on 4-12% SDS-PAGE gel. C) Isothermal titration calorimetry of MBP-ATG9A (830-839) and ATG13 ^HORMA^:TSF-ATG101. D) GST pulldown assay using purified TSF-ATG13^HORMA^:GST-ATG101 incubated with lysate containing ATG9A wild type or mutants. Bound protein was run on 4-12% SDS-PAGE gel. E) Quantification of GST pulldown assays from D) showing standard deviation (n=3).

ATG9A is the only known ubiquitously expressed transmembrane protein of the autophagy initiation cascade in mammals (Young et al. 2006). ATG9A forms Golgi-derived vesicles which are recruited to autophagy initiation sites through a complex trafficking process (Claude-Taupin et al. 2021; Davies et al. 2018; Judith et al. 2019; Lamb et al. 2016; Mattera et al. 2017). Global knock out of *ATG9A* in mice results in impaired autophagosome biogenesis and accumulation of p62/SQSTM1 aggregates (Saitoh et al. 2009), and conditional knockout in the brain leads to progressive neurodegeneration (Yamaguchi et al. 2018). Structurally, ATG9A is composed of two distinct domains, a transmembrane domain (TMD, 1-525) and a C-terminal intrinsically disorder region (IDR, 526-839) (Figure 1A). The TMD of ATG9A forms an interlocked trimer and functions as a lipid scramblase (Guardia et al. 2020; Maeda et al. 2020; Matoba et al. 2020). ATG9A distributes incoming ER synthesized lipids, transported via ATG2 proteins, across the growing phagophore from the outer to inner leaflet (Gómez-Sánchez et al. 2018; Schü et al. 2020). The IDR region of ATG9A contains sites of regulatory signaling including phosphorylation via TBK1 and ULK1, ubiquitination, and direct protein-protein interactions (Wang et al. 2022; Weerasekara et al. 2014; C. Zhou et al. 2016). It has been shown that yeast Atg9 is capable of acting as a seed for phagophore initiation (Sawa-Makarska et al. 2020) and it seems reasonable to expect that mammalian ATG9A might do the same. To carry out any of their lipid transfer, regulatory, assembly, or putative seeding functions, ATG9A vesicles must first be recruited to sites of autophagy initiation, the focus of the present study.

The ATG9A, ATG13 and ATG101 form a multifunctional hub at an early stage of starvation-induced autophagy and mitophagy initiation (Dalle Pezze et al. 2021; Karanasios et al. 2016; Zachari et al. 2019). A BioID mass spectroscopy approach revealed that ATG13:ATG101 can recruit ATG9A during p62-dependent autophagy, independent of FIP200 and ULK1 (Kannangara et al. 2021). Deletion of ATG13 or ATG101 led to a mislocalization of ATG9A, leading in turn to an accumulation of p62 aggregates identical to the ATG9A KO phenotype (Kannangara et al. 2021; Saitoh et al. 2009). This study highlighted the importance of the ATG9A, ATG13 and ATG101 nexus and focused our attention on this subnode of the autophagy interaction network. Clearance of damaged mitochondria requires both the ULK1 complex and ATG9A to promote efficient mitophagy (Itakura et al. 2012). During PINK1/Parkin dependent mitophagy the ubiquitination of outer mitochondrial membrane proteins promotes recruitment of OPTN, NDP52 and p62 (Heo et al. 2015; Lazarou et al. 2015; Wong and Holzbaur 2014). However, TBK1-phoshorylated OPTN has been reported to directly recruit ATG9A during mitophagy, thereby bridging cargo to ATG9A vesicles in manner that could potentially bypass the need for ATG13 to recruit ATG9A (Yamano et al. 2020; Z. Zhou et al. 2021). This contrasts with the strict dependence of NDP52-mediated selective autophagy on the ULK1 complex for initiation, which led us to focus on the role of the ATG9A:ATG13:ATG101 complex in NDP52-dependent PINK1/Parkin mitophagy. Here, we have identified the binding interface between ATG9A, ATG13, and ATG101. We term the region of the extreme C-terminus of ATG9A that binds to the ATG13:ATG101 HORMA dimer the “HORMA dimer interacting region” (HDIR). We determined the structure of the human ATG9A HDIR bound to the ATG13:ATG101 HORMA dimer and confirmed that the interface functions in NDP52 dependent mitophagy.

## Results

### Mapping the ATG9A region that interacts with the ATG13:ATG101 HORMA dimer

To investigate the interaction between ATG9A and the ULK1 complex, ATG9A, ATG13, and ATG101 proteins were overexpressed and purified via mammalian HEK293GnTi cells. The HORMA domain (1-197) of ATG13 (henceforth called ATG13^HORMA^) was purified along with ATG101. An amylose bead pulldown assay using full length and N-terminal truncations of ATG9A as bait to recruit ATG13^HORMA^:ATG101 was performed (Figure 1B). Full length MBP-ATG9A in detergent micelles strongly bound ATG13^HORMA^:ATG101. To study the isolated soluble C-terminal domain of ATG9A as a trimeric assembly, we engineered a FOLDON domain N-terminal to the ATG9A C-terminal constructs. FOLDON is a small 8 kDa N-terminal domain of T4 fibritin which intrinsically forms trimers (Meier et al. 2004). MBP-FOLDON-ATG9A CTD constructs containing 692-839, 723-839, 801-839, 828-839, and 830-839 all recruited ATG13^HORMA^:ATG101 (Figure 1B). These data show the last nine residues of ATG9A (termed the ATG9A tail) are sufficient to recruit ATG13 ^HORMA^:ATG101 (Figure 1B). Monomeric MBP-ATG9A (692-839) also recruited the ATG13 ^HORMA^:ATG101, therefore the trimeric assembly is not required. We designate human ATG9A residues 830-839 as the “HORMA dimer interacting region” (HDIR).

To determine the affinity of the ATG9A HDIR for ATG13^HORMA^:ATG101 we used isothermal titration calorimetry (ITC). Serial injection of MBP-ATG9A (801-839) into a cell containing ATG13^HORMA^:ATG101 resulted in a *K*_d_ of 2.4 ± 0.3 mM (Figure 1C). This assay showed equal stoichiometry between the ATG13^HORMA^:ATG101 dimer and ATG9A 801-839 (N=0.82 ± 0.03) (Figure 1C). This ratio is consistent with the pulldown result that the trimeric assembly is not required for efficient binding. An alanine scan of the ATG9A tail was performed to determine the effect of tail residue side chains on (801-839):ATG101 binding. Recruitment of wild type MBP-ATG9A, V839A, V836A, Q835A, L832A, and a double mutant (DE830-831AA) was robust in the presence of ATG13^HORMA^:ATG101 (Figure 1D, E). Mutants K838A and P833A reduced the binding efficiency to half when compared to wild type. The mutations H837A and P834A abolished the binding of ATG13^HORMA^:ATG101, reducing it to 20% and 5% of wild type, respectively. Truncation of residues 830-839 from the entire ATG9A tail similarly abolished the interaction. These data identified the ATG9A tail (830-839) as the site of direct interaction between ATG13^HORMA^:ATG101, linking ATG9A to the ULK1 complex.

### Crystal Structure of the ATG9A HDIR:ATG13^HORMA^:ATG101 Ternary Complex

Having mapped the HDIR, we pursued structural analysis of the ternary complex. Multisequence alignment showed that the residues within the HDIR are conserved across the Opisthokonta, with the exception being *S. cerevisiae*, whose Atg9 ortholog has no sequence homology with the human HDIR (Figure 2A). P834 and H837 are highly conserved across Opisthokonta ATG9A HDIR sequences, excluding budding yeasts. Alphafold2 predictive modeling (Jumper et al. 2021) was performed using the primary sequence of the ATG13^HORMA^, ATG101, and the ATG9A tail residues (828-839). To generate a set of models the primary amino acid sequences of ATG9A, ATG13^HORMA^, and ATG101 were submitted in varying input positions. Thirty models were generated which were overlayed to determine the position of the HDIR (Supp Fig 2A). Two distinct positions were observed, one that involved binding only to ATG13^HORMA^ and one which spanned both HORMA domains. The models with the HDIR at the dimer interface had higher pLDDT scores. From these models, the HDIR was located adjacent to the N-terminus of ATG101. We designed a fusion construct between the HDIR and ATG101, using a flexible linker of 5 residues (GSDEA) to increase the affinity of the ternary complex for crystallization (Figure 2B).

**Figure 2.**
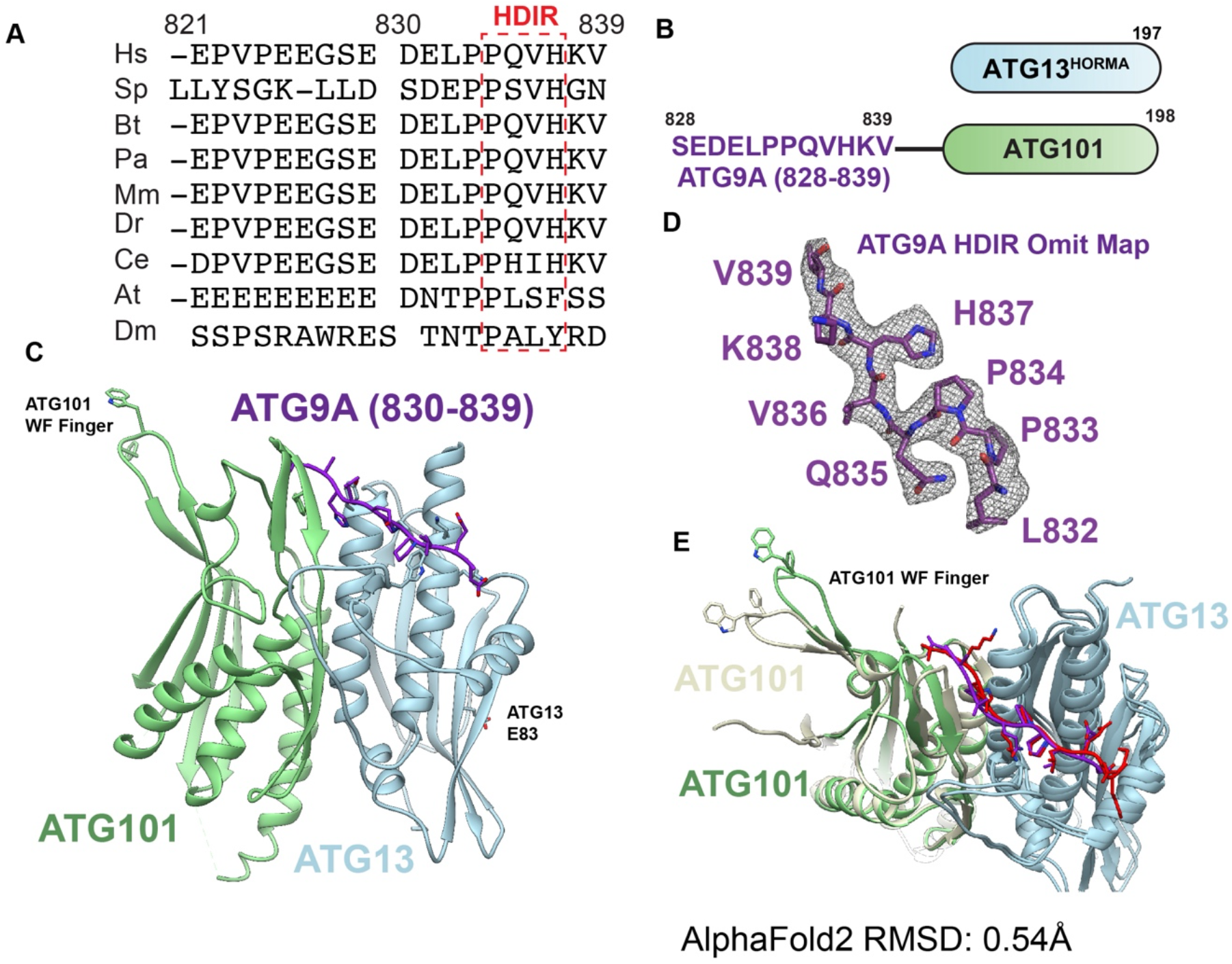
Ternary ATG9A HDIR:ATG13 ^HORMA^:ATG101 Crystal Structure. A) Multisequence alignment of ATG9A C-terminal tails performed using ClustalO and aligned to ATG9A from Hs. Hs = Homo Sapiens, Sp=, Bt=, Pa=, Mm=, Dr=, Ce=, At=, Dm= B) Constructs used for crystallization including, ATG13 ^HORMA^ (1-197) and fused ATG9A (829-839)-GlySerLinker-ATG101 (1-198) C) Ribbon representation of the ATG9A HDIR-ATG101 and ATG13 ^HORMA^ constructs for crystallography colored by protein. ATG13 ^HORMA^ -light blue, ATG101-light green, ATG9A – purple D) Omit map of the ATG9A HDIR shown at 2.9s E) Overlay of crystal structure and Alphafold2 predicted model aligned to the ATG13 ^HORMA^ domain. Crystal structure colored as in (C), Alphafold2 model colored as ATG13 ^HORMA^ - light blue, ATG101 – yellow and ATG9A tail – red.

We determined the crystal structure of the ATG9A HDIR-ATG101 fusion construct bound to ATG13^HORMA^ to 2.4 Å resolution (Figure 2C, Table 1). Molecular replacement was performed using the ATG13^HORMA^:ATG101 apo structure (PDB:5C50) (Qi et al. 2015). The asymmetric unit contains two ATG9A HDIR: ATG13^HORMA^:ATG101 complexes, both in a similar conformation, except that the ATG101 WF finger is in different conformations in the two complexes, due to differences in the crystal packing environment. ATG13^HORMA^ and ATG101 are in essentially the same conformation as in the apo ATG13^HORMA^:ATG101 structure with a root mean square deviation (RMSD) of 0.45 Å across all backbone atoms (Supp Figure 2B-E). Electron density at the interface for ATG13^HORMA^:ATG101 fit all ten residues of the HDIR, allowing confident positioning of residues 830-839 (Figure 2D). The ATG13^HORMA^:ATG9A and ATG101:ATG9A interfaces bury 388 Å^2^ and 241 Å^2^ of solvent-accessible surface area, respectively. Overall, our experimental structure agrees with the Alphafold2 prediction with an RMSD of 0.54 Å for backbone atoms (Figure 2E). Our ternary crystal structure of ATG9A HDIR: ATG13^HORMA^:ATG101 complex determined that the HDIR binds at the interface between ATG13 and ATG101 HORMA domains.

**Table 1.** Data collection and refinement statistics.

|  | ATG9 HDIR-ATG101:ATG13 Complex |
| --- | --- |
| **Data collection** |  |
| Space group | P 2<sub>1</sub> 2<sub>1</sub> 2<sub>1</sub> |
| Cell dimensions |  |
| *a*, *b*, *c* (Å) | 45.45, 139.86, 147.60 |
| $\alpha$, $\beta$, $\gamma$ (°) | 90, 90, 90 |
| Resolution (Å) | 39.41-2.41 (2.496-2.41) |
| *R*<sub>pim</sub> | 0.03972 (0.5303) |
| *I* / σ*I* | 14.55 (1.31) |
| *CC*<sub>1/2</sub> | 0.999 (0.559) |
| Completeness (%) | 99.75 (99.89) |
| Redundancy | 6.6 (6.8) |
| **Refinement** |  |
| Resolution (Å) | 39.41-2.41 |
| No. reflections | 37256 (3659) |
| *R*<sub>work</sub> / *R*<sub>free</sub> (%) | 21.3 (29.4) / 26.1 (35.6) |
| No. atoms | 6307 |
| Protein | 5867 |
| Ligand/ion | 252 |
| Water | 188 |
| *B*-factors | 64.5 |
| Protein | 63.44 |
| Ligand/ion | 89.94 |
| Water | 64.17 |
| R.m.s. deviations |  |
| Bond lengths (Å) | 0.005 |
| Bond angles (°) | 0.70 |
| Clash score | 5.56 |
| Ramachandran plot |  |
| Favored (%) | 96.95 |
| Allowed (%) | 2.52 |
| Outliers (%) | 0.53 |

Values in parentheses are for highest-resolution shell.

### ATG13^HORMA^ and ATG101 Form a Hydrophobic Groove for ATG9A HDIR Binding

The interaction of the HDIR with the ATG13^HORMA^:ATG101 dimer is mediated by backbone and sidechain interactions. The loop between β2’ and αB in ATG13^HORMA^ (T46-W50) is positioned to support the binding of the HDIR (Figure 3A). This rearrangement supports a hydrogen bonding interaction between ATG9A L832 and Q835 and ATG13 K15 and G47 (Figure 3A). The loop region of G43-Y45 in ATG101 joins a β-sheet network between β2 and HDIR residues H837-V839 (Figure 3B). We termed this new β-sheet, β1’ (SuppFig 2E). These two conformational changes in ATG13^HORMA^:ATG101 occur in the presence of ATG9A binding and are not observed in apo ATG13^HORMA^:ATG101 structures. The HDIR peptide binds across both HORMA domains within a long hydrophobic groove (Figure 3C). ATG101 β2’ (Y45) and ATG13 αA (K15, F16, K18, F19), αA-B connector (W50) and αC (Y115, Y118) form a greasy patch of residues which support the full length of the HDIR.

**Figure 3.**
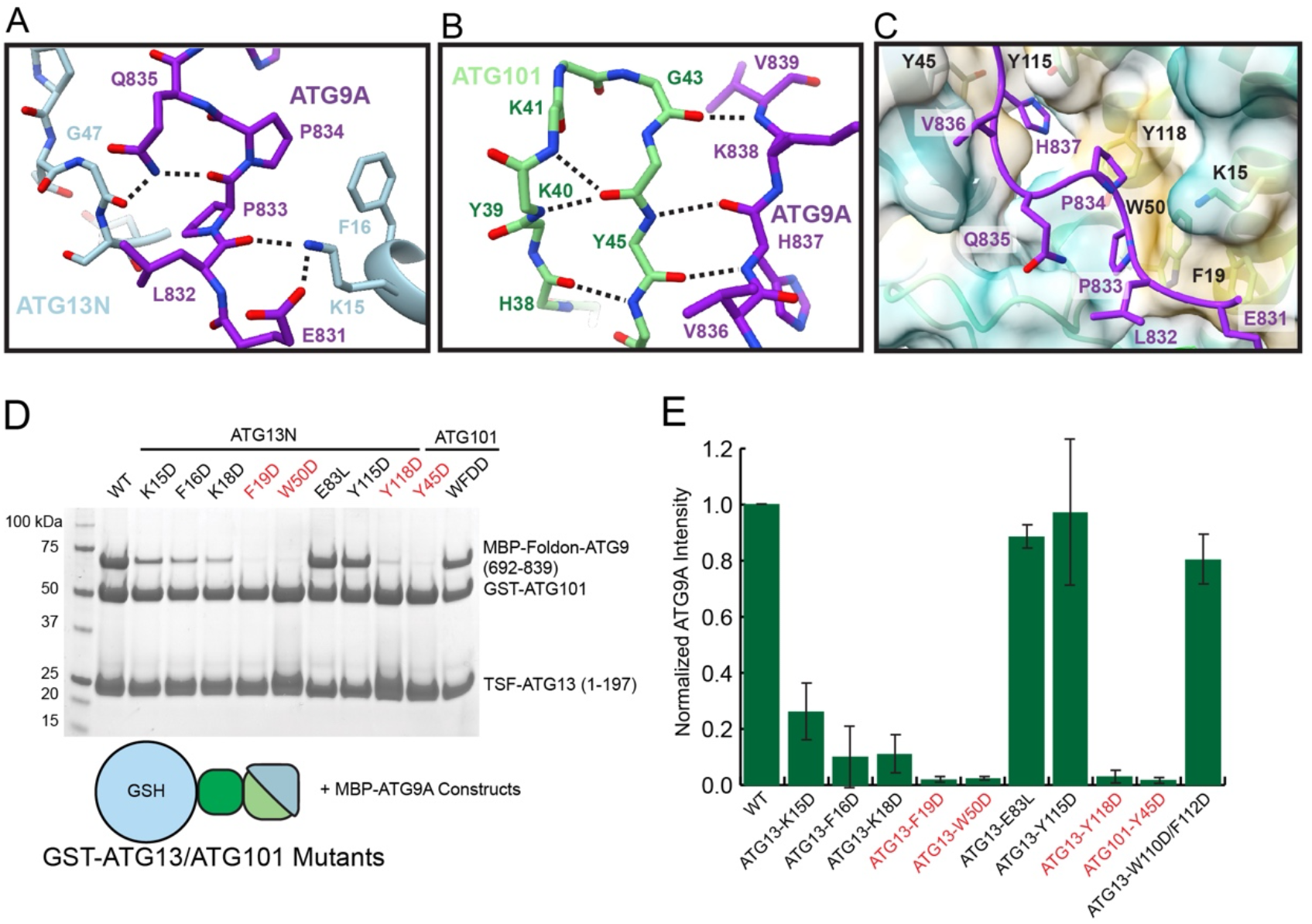
Interfaces Between ATG9A HIDR and ATG13 ^HORMA^:ATG101. A) Close up of the hydrogen network between ATG13 ^HORMA^ and the ATG9A HDIR. B) The extended β-sheet between ATG101 and ATG9A C) View of the hydrophobic groove across the HORMA dimer interface, colored by hydrophobicity D) GST pulldown assay using purified wild type TSF-ATG13 ^HORMA^:GST-ATG101 and listed mutants incubated with lysate containing MBP-Foldon-ATG9A. Position of mutants E83 and WF finger shown in Figure 2C. E) Quantification of GST pulldown assays from D) showing standard deviation (n=3).

To validate the ATG13^HORMA^: ATG101:HDIR interface, we used a GST pulldown assay to monitor the recruitment of ATG9A (692-839) to wild type and mutated ATG13^HORMA^:ATG101 proteins. Wildtype ATG13^HORMA^:ATG101 robustly recruited ATG9A (Figure 3D, E). Mutations of hydrophobic residues in ATG13^HORMA^ (K15D, F16D, K18D, Y115D) severely impaired ATG9A binding. Removing the hydrophobic residues, F19D, W50D, Y118D in ATG13 ^HORMA^ and Y45D in ATG101 fully abolished binding of ATG9A. Mutation E83L in ATG13^HORMA^, which is homologous to the binding site of budding yeast Atg13 to Atg9, had no effect on binding (S. W. Suzuki et al. 2015). Removing the WF finger of ATG101 (WF to DD) had no effect on ATG9A binding. Our mutational analysis suggests that the hydrophobic groove drives ATG9A binding and explains the need for both ATG101 and ATG13 in binding ATG9A in vivo (Kannangara et al. 2021).

### ATG9A HDIR binding to ATG13^HORMA^:ATG101 facilitates NDP52-dependent mitophagy initiation

We sought to determine if the HDIR interaction with the HORMA dimer plays a role in PINK1/Parkin mitophagy initiation. We focused on NPD52-mediated mitophagy because of the evidence that OPTN can directly bind and recruit ATG9A (Yamano et al. 2020), whereas NDP52 relies on direct binding to the ULK1 complex (Ravenhill et al. 2019; Vargas et al. 2019) to mediate ATG9A recruitment. The *ATG13* gene was deleted in the context of the mitophagy receptor penta-KO (OPTN, NDP52, TAX1BP1, p62 and NBR1) cell line (SuppFig. 2) (Lazarou et al. 2015). Turnover of the inner mitochondrial protein cytochrome C oxidase subunit II (COXII) during mitochondrial stress conditions was monitored to assay mitophagy. In the absence of NDP52 and ATG13, mitochondrial degradation was impaired (Figure 4A,B) as shown by the robust COXII signal after 20 hours of oligomycin-antimycin treatment to induce mitochondrial depolarization and mitophagy. Rescue experiments with NDP52 alone were insufficient to restore COXII degradation defects in ATG13 KO/penta KO cells (Figure 4A,B). COXII degradation was increased when both wild type ATG13 and NDP52 were added. Rescue experiments using ATG13 W50D and Y118D mutations, which disrupt the interaction between ATG13 and ATG9A in vitro (Figure 3C), showed reduced recovery of COXII degradation when compared to rescue with wild type ATG13 (Figure 4C). These findings demonstrate that the HDIR binding interface on ATG13 contributes to NDP52 dependent PINK1/Parkin mitophagy.

**Figure 4.**
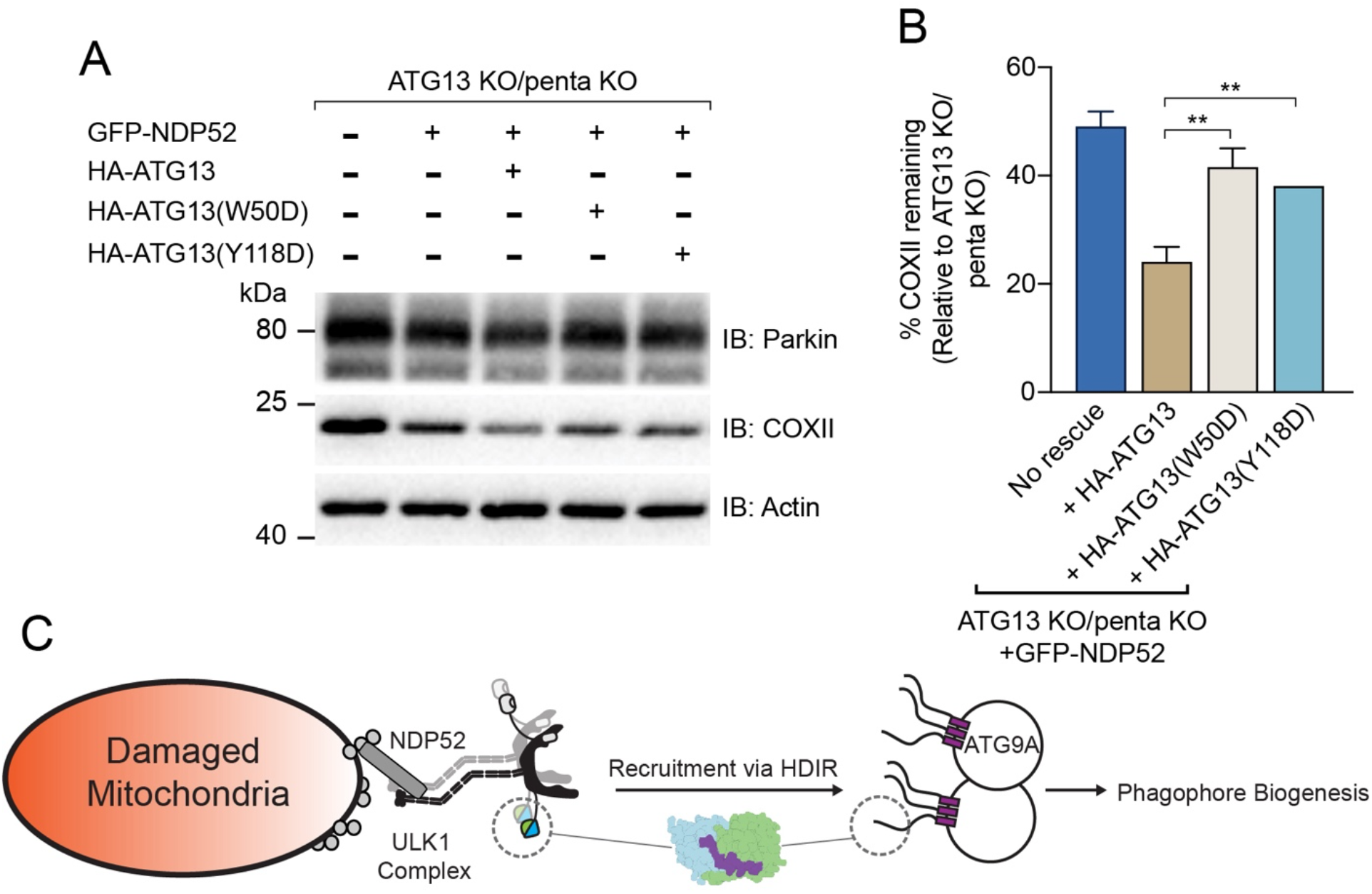
Rescue assays of COXII Degradation in ATG13 KO/penta KO cells and model for ATG9:ATG13:ATG101 mediated mitophagy initiation. A) ATG13 KO/penta KO expressing BFP-Parkin along with GFP-NDP52 and ATG13 proteins as indicated. Protein levels analyzed by immunoblotting after 20 hours of OA treatment along with B) quantification of COXII levels (n=2) C) Model schematic of ATG9:ATG13:ATG101 interaction in NDP52 mitophagy

## Discussion

Understanding the interactions responsible for autophagy initiation is central to efforts to therapeutically modulate bulk and selective autophagy pathways and to understand their functioning at a fundamental level. Here, we determined the structure of the ATG9A HDIR bound to the ATG13:ATG101 HORMA dimer (Figure 2C). As a central part of the autophagy initiating ULK1 complex, the HORMA dimer is considered a key element enabling the ULK1 complex to recruit ATG9A vesicles for autophagy initiation (Dalle Pezze et al. 2021; Karanasios et al. 2016; Zachari et al. 2019). The structure helps bring an understanding of ATG9A binding in the context of other structural information on the HORMA dimer, and HORMA domains in general. The HDIR site overlaps a hydrophobic binding site for the small molecule benzamidine that was identified in the apo structure of the human HORMA dimer (Qi et al. 2015), explaining the normal function of this site. Other HORMA domain proteins function as protein interaction hubs regulated by conformational changes in the safety belt region (Gu, Desai, and Corbett 2022). This posed the question as to whether interactions of the ATG13:ATG101 dimer could be regulated by conformational switching as seen for Mad2 and other HORMA domains (Gu, Desai, and Corbett 2022). The interaction between the ATG9A HDIR and ATG13:ATG101 is distal to and thus apparently independent of the safety belt region.

Our biochemical experiments confirmed that the ATG13 residues that bind the ATG9A HDIR in the structure are required for full function. We noticed that none of these mutations completely abolishes mitophagy (Figure 4A,B). On the other hand, the deletion of the entire ATG13 HORMA domain has a nearly complete loss of function, as seen by Kananngara et al. (2021). Thus, the HORMA dimer has additional functions distinct from the ATG9A tail binding pocket. We confirmed that the essential WF motif of ATG101 (H. Suzuki et al. 2015) is not involved in ATG9A binding. An intact ATG13 HORMA dimer is required for ATG101 to associate with the rest of the ULK1 complex via ATG13 IDR (Qi et al. 2015). The unaccounted-for residual function of the HORMA dimer seems likely related to the role of the WF finger, which will be important to establish.

Human ATG9A contains a long C-terminal IDR, whereas budding yeast Atg9 contains an extensive N-terminal IDR. Sequence alignment of the ATG9A tail shows a strong conservation across most of the Opisthokonta (Figure 2A) except budding yeast, which lacks the HDIR. *S. cerevisiae*, as well as related thermophilic yeasts such as *K. lactis* and *L. thermotolerans* that have been used as model systems in autophagy, has distinct initiation machinery when compared to other Opisthokonta (Hurley and Young 2017). The ULK1 (human) kinase complexes contains ATG13 along with its constituent binding partner ATG101. However, budding yeast entirely lacks an ATG101 ortholog, and its functions are apparently replaced by the single HORMA domain in yeast Atg13 (Jao et al. 2013). Thus, even though budding yeast Atg9 binds to the HORMA domain of Atg13 (S. W. Suzuki et al. 2015), the mammalian sequence motif and structural interactions do not seem to exist in yeast.

We combined biochemical analysis and predictive modeling to generate a fusion ATG9A-ATG101 protein for experimental structure determination. Protein structure prediction has rapidly advanced via deep learning algorithms such as Alphafold2 (AF2) (Baek et al. 2021; Jumper et al. 2021). Predictive modeling can determine the binding pockets of polypeptide chains with < 2 Å RMSD (Ko and Lee 2021). However, validation of structural models remains a critical step in guiding our understanding of biological mechanisms. Minimizing the amount of primary sequence during modeling can greatly improve the accuracy of predicted interfaces. Our truncation data limited the primary sequence for the ATG9A component to be modeled on the ATG13:ATG101 dimer, making AF2 prediction of the complex feasible. Importantly, determining which of the two modeled binding sites was correct relied on experimental information. The potential for predictive models combined with in vitro biochemical analysis to determine protein-protein interfaces correctly will continue to grow as more methods are explored and structural studies are guided by predictive models. In our study, we found that experimental interaction mapping provided essential constraints prior to AF2 prediction of the complex, and downstream model validation by mutational and structural analysis was essential downstream.

Ubiquitinated autophagy cargoes recruit receptors (OPTN, p62, NBR1, NDP52, TAX1BP1) and some bind directly to the ULK1 complex via FIP200 (Ravenhill et al. 2019; Turco et al. 2019; Vargas et al. 2019). This includes NDP52 recognition of Parkin ubiquitination marks on mitochondria damaged by uncoupling agents (Vargas et al. 2019). These interactions drive clustering and allosteric activation (Shi, Chang, et al. 2020) of the ULK1 complex to promote the autophagy initiation cascade. While the micromolar affinity of the ATG9A HDIR:ATG13:ATG101 interaction is moderate, it is easy to see how the clustering of multiple ULK1 complexes on an extended ubiquitinated platform such as a damaged mitochondrion would contribute avidity to recruitment of ATG9A trimers. The 1:1 stoichiometry established here implies that a single ATG9A trimer can bind to three ULK1 complexes presented in a cluster on a cargo substrate (Figure 4C). The interaction between the C-terminal IDRs of the ATG13 molecules and the N-terminal crescent domains of the FIP200 molecules would position the massive C-terminal coiled coil domains of FIP200 distal to the ATG9A vesicles. Based on the dimensions of FIP200 (Shi, Yokom, et al. 2020), this would allow for fly-casting for ATG9A vesicles from roughly 35 nm from sites of clustered ubiquitin.

While our model presents a satisfying and plausible model for ATG9A vesicle recruitment downstream of the ULK1 complex in autophagy initiation, this is almost certainly not the complete mechanism, nor the only mechanism. Kananngara (2021) found that ULK1 could still interact with ATG9A in the absence of ATG13 as measured by BioID, although this was insufficient to support normal levels of p62 autophagy. Thus, at least one additional point of contact between ATG9A and the ULK1 complex is likely to exist, at least in some forms of autophagy. A more significant departure from this mechanism is suggested by the finding that OPTN can recruit ATG9A directly, bypassing the need for the ATG9A-binding role of the ULK1 complex (Yamano et al. 2020). The physiological rationale for these various pathways remains largely unknown, but the distinct role of different cargo receptors in neurodegeneration suggests it will be important to better understand. Elucidating the molecular details of distinct initiation mechanisms in various “flavors” of mitophagy should provide windows into disease mechanism, and ultimately therapy.

## Materials and Methods

### Plasmid construction

Plasmids were engineer and modified according to the published protocol (dx.doi.org/10.17504/protocols.io.bxrjpm4n). The fusion construct of Human ATG9A C-terminal with ATG101 for crystallization was subcloned as an N-terminal GST tag, TEV cleavage site, ATG9 (828-839), 5 aa linker (GSDEA) followed by ATG101 (1-198) into the pCAG vector. ATG13 (1-197) (no tag) was subcloned into the pCAG vector using ClaI/XhoI sites. The ATG101:ATG13 HORMA construct for Isothermal titration calorimetry (ITC) experiment was described (Qi, et al., 2015). Proteins were tagged with GST, MBP or TwinStrep-Flag (TSF) for affinity purification or pull-down assays. All constructs were verified by DNA sequencing. Details are shown in the key resources table and plasmids are deposited to Addgene.org (Addgene ID = 188073-188106).

### Protein expression and purification

For crystallization, GST-ATG9 (828-839)-ATG101 were co-transfected with ATG13 (1-197) using the polyethylenimine (PEI)-MAX (Polysciences) transfection system. Cells were transfected at a concentration of 2 × 10^6^/ml. After 48 hours, cells were pelleted at 500x g for 10 min, washed with PBS once, and then stored at −80°C. The pellets were lysed with lysis buffer (25 mM HEPES pH 7.5, 200 mM NaCl, 2 mM MgCl_2_, 1 mM TCEP, 5 mM EDTA, 10% Glycerol) with 1% Triton X-100 and protease inhibitor cocktail (Thermo Scientific, Waltham, MA) before being cleared at 17000x rpm for 35 min at 4°C. The clarified supernatant was purified on GST Sepharose 4B resin (GE healthcare) and then eluted in the lysis buffer with 25mM glutathione. A general protocol for purification of GST-tagger ATG13:ATG101 constructs has been uploaded online (dx.doi.org/10.17504/protocols.io.n92ldzx59v5b/v1). After His_6_-TEV cleavage at 4°C overnight, the sample was diluted 5 times in SP column buffer A: 30 mM MES pH 6.0, 3 mM b-mercaptoethanol (BME) and then loaded onto a HiTrap SP HP 5 ml column (GE healthcare, Piscataway, NJ). Elution from the SP column was performed with a 70 ml linear gradient from 0–500 mM NaCl in SP buffer A. The cleavage sample was eluted at the buffer conductivity of ∼ 25 mS/cm. After each fraction was analyzed by SDS gel, the pooled fractions were concentrated in Amicon Ultra15 concentrator (MilliporeSigma, Burlington, MA) and exchanged buffer to the ITC buffer (25 mM HEPES pH 7.5, 150mM NaCl, 1mM MgCl2, 1mM TCEP).

For ITC experiment, GST-ATG13 (12-200):ATG101 (1-198) complex was expressed in *Spodoptera frugiperda* (Sf9) cells. Baculoviruses were generated in Sf9 cells with the bac-to-bac system (Life Technologies). Cells were infected and harvested after 72 hr. Purification was performed as described above for HEK cell expression.

MBP-ATG9A related constructs were transfected in HEK GnTi-cells at 2× 10^6^ /ml density. Cells were harvested after 48 hr transfection. For ATG9A CTD constructs, the pellets were lysed with lysis buffer/protease inhibitor cocktail/1% Triton X-100 at room temperature for 15min. The clarified supernatant was purified on amylose resin (New England biolabs, Ipswich, MA) and then eluted in the lysis buffer with 40 mM Maltose (Sigma, St Louis, MA). The eluted samples were further concentrated then loaded onto a Superose 6 Increase 10/300 GL column (Cytiva, Marlborough, MA) in ITC buffer. For MBP-ATG9A full protein, cells were lysed in lysis buffer/protease inhibitor cocktail/1% DDM/CHS. All wash, elution or pull-down buffers contain 0.05% DDM/CHS. Purification of ATG9A truncations via Amylose resin has been deposited online (dx.doi.org/10.17504/protocols.io.6qpvr6nkovmk/v1)

TSF-ATG101:ATG13 (1-197) complex used for pull-down assay was expressed in HEK GnTi-cells at 2× 10^6^ /ml density. After 48 hr transfection, the pellets were lysed with the lysis buffer/1% Triton X-100 at room temperature for 15min. The clarified supernatant was purified on strep-tactin resin (IBA Lifesciences, Germany). After extensive wash, the sample was eluted with lysis buffer/4 mM desthiobiotin, then loaded onto a Superdex 200 Increase 10/300 GL column (Cytiva) in ITC buffer. Method for purification of Twin-STREP FLAG tagged ATG13:ATG101 dimers has been uploaded online (dx.doi.org/10.17504/protocols.io.yxmvmn4yng3p/v1)

### Bead pull down assays

GST-ATG101:TSF-ATG13 (1-197) HORMA WT or mutants were expressed in 10 ml of HEK-GnTi cells. The cells were harvested 48 hours after transfection. The pellets were homogenized in 0.5 ml of lysis buffer/protease inhibitors/1% TritonX-100, and clarified after 40,000g x 15 min. The lysate was incubated with 30 µl Glutathione Sepharose beads (GE Healthcare) at 4°C for 4 hours. The beads were washed twice by lysis buffer, then incubate with recombinant MBP proteins (final 2 µM) or lysate from 10 ml HEK cells at 4°C overnight. Next day, the beads were washed four times, and then elute in 50 µl ITC buffer/25mM glutathione. 17 μl eluent was mixed lithium dodecylsulfate (LDS)/BME buffer, heated at 60°C for 5 min and subjected to SDS/PAGE gel. Protocol for GST resin based pulldown assay is deposited online (dx.doi.org/10.17504/protocols.io.kqdg3pdrpl25/v1)

In 300 µl, recombinant MBP proteins (final 1.2 µM), TSF-ATG101:ATG13 (1-197) complex (final 3 μM) were incubated with 30 μl Amylose resin (New England Biolabs, Ipswich, MA) at 4 °C overnight in the ITC buffer. The beads were washed 4 times, and then eluted in 50 μl ITC buffer/50 mM Maltose. 17 μl eluent was mixed lithium dodecylsulfate (LDS)/BME buffer, heated at 60°C for 5 min and subjected to SDS/PAGE gel. Band intensities were quantified using Fiji ImageJ2 (version 2.3.0/1.53f). Raw data from this assay has been uploaded to Zenodo (10.5281/zenodo.68095410) and protocol published online (dx.doi.org/10.17504/protocols.io.e6nvwk7xwvmk/v1).

### Isothermal titration calorimetry

All samples were dialyzed against the ITC buffer at 4 C overnight. The sample cell contained 0.2 ml of 17 μM ATG101-ATG13 HORMA complex, and MBP-ATG9 (801-839) (240 μM) was added in 13 injections of 2.7 μl each. Measurements were repeated four times and carried out on a MicroCal PEAQ-ITC 1.4.1 instrument according to the manual (Malvern Panalytical Inc, Westborough, MA). The data were processed using MicroCal PEAQ-ITC analysis software. The binding constant (*K*_d_) was fitted using a one-site model.

### Modeling, Crystallization, and crystallographic analysis

Alphafold2 was run using the ColabFold notebook (https://github.com/sokrypton/ColabFold) using version 2.1 on default settings. Mutlisequence alignment of Figure 2A was performed in seaview version 5.0.4 (Gouy, Guindon, and Gascuel 2010). The ATG9 HDIR (828-839) fused ATG101 (1-198):ATG13 (1-197) complex was concentrated to 6 mg/ml in the ITC buffer. Crystallization was carried out by sitting-drop vapor diffusion using an automated liquid-handling system (Mosquito, TTP LabTech, UK) at 288 K in 96-well plates. The protein solution was mixed with the reservoir buffer composed of 0.1 M HEPES pH 7.5, 0.2 M NaCl, 12% PEG8000 with a ratio of 1:1. The crystal was obtained in 2–4 days. Crystals were cryo-protected in 28% glycerol/reservoir buffer and frozen in liquid N_2_. Protocol for purification of ATG9 HDIR-ATG101:ATG13^HORMA^ complex and crystallization has been deposited (dx.doi.org/10.17504/protocols.io.dm6gpbkdjlzp/v1)

Native data were collected from a single frozen crystal using a Dectris Pilatus 6M detector at beamline 12-2, SSRL. All data were processed and scaled using XDS 4.0 (Kabsch, 2010). The crystal diffracted to 2.4 Å resolution and belonged to space group P2_1_2_1_2_1_ with unit cell dimensions a = 45.453 Å, b = 139.86 Å, c = 147.595 Å, α = β = γ = 90°. A molecular replacement solution was found using partial structures derived from ATG101:ATG13 HORMA apo structure (PDB: 5C50) as a search model with Phenix (version 1.20.1-4487) (Adams et al. 2010). Model building and refinement were carried out using Coot 0.9.6 EL (Emsley et al. 2010) and Phenix (version 1.20.1-4487) (Adams et al. 2010). Structural figures were generated with PyMol (version 2.5) (Schrödinger 2018) or UCSF Chimera (version 1.16) (Pettersen et al. 2004).

### Generation of *ATG13* KO/penta KO using CRISPR/Cas9

HeLa cells (ATCC; CCL-2.2) were maintained in DMEM with 10% FBS, 4.5 g/l Glucose (Sigma, G8769), 1x GlutaMAX^™^ (ThermoFisher, 35050061), 1x MEM NEAA (ThermoFisher, 11140-050), 25 mM HEPES (1688449). Penta KO cells missing five autophagy adaptor proteins (p62, OPTN, NDP52 TAX1BP1 and NBR1) were described previously (Lazarou et al. 2015). CRISPR guide RNA (gRNA) that targets *ATG13* (TTGCTTCATGTGTAACCTCTGGG) was cloned into pSpCas9(BB)-2A-GFP vector (a gift from Feng Zhang; Addgene plasmid # 48138) via Gibson Cloning kit (New England Biolabs). The construct was then sequenced and transfected into penta KO cells with X-tremeGENE 9 (Roche) overnight and GFP positive single cells were sorted by fluorescence activated cell sorting (FACS) into 96 well plates. Single cell colonies were then screened by immunoblotting with an anti-ATG13 antibody (Cell Signaling; 13468, RRID: AB_330288). Gene editing in the positive clones was identified by Sanger sequencing. Protocol for creating this cell line as well as the cell line itself are in progress of being deposited.

### CoxII degradation assay to assess mitophagy

Mitophagy treatment was previously described (DOI: 10.1038/s41467-019-08335-6; DOI: 10.1016/j.molcel.2021.03.001). Briefly, 350,000 cells were seeded the day before the treatment day in 6 well plates. Rescue experiments were performed with eGFP-NDP52 (Addgene:188785), HA-ATG13 (Addgene:186223), HA-ATG13(W50D) (Addgene:188639), HA-ATG13(Y118D) (Addgene:188638). The cells were then incubated with growth media containing 4 μM Antimycin A, 10 μM Oligomycin and 10 μM QVD for 20 h. Following treatment, the cells were washed with ice-cold 1x PBS, harvested using cell scrapers and lysed in lysis buffer containing 1× LDS sample buffer (Life Technologies) and freshly added 100 mM dithiothreitol (DTT; Sigma). Samples were heated at 99 °C with shaking for 7 min. Approximately 25μg of protein per sample was run on 4-12% Bis-Tris gels (Life Technologies) according to manufacturer’s instructions. Gels were electro-transferred to polyvinyl difluoride membranes (PVDF) and immunoblotted with antibodies against Actin (Cell Signaling; 4967S, RRID: AB_330288), Parkin (Santa Cruz; sc-271478, RRID: AB_628104), COXII (Abcam; ab110258, RRID: AB_10887758). For western blot quantification, band intensities were measured with ImageLab 5.2.1 (BioRad). Statistical significance was calculated from two independent experiments using one-way ANOVA. Error bars are means ± standard deviation. Raw data from this assay has been uploaded to Zenodo (10.5281/zenodo.68095410) and protocol for this assay is in progress to be deposited at protocol.io.

## Data Availability

Coordinates and structure factors have been deposited in the Protein Data Bank under accession code PDB 8DO8. Protocols have been deposited in protocols.io. Plasmids developed for this study will be deposited at Addgene.org. Raw data files for gel scans have been uploaded to Zenodo (10.5281/zenodo.68095410).

## ACKNOWLEDGEMENTS

We thank members of the Hurley lab, Dorotea Fracchiolla, and others in Aligning Science Across Parkinson’s Team mito911 for advice and discussions. We thank Clyde Smith and Lisa Dunn at SSRL beamline BL12-2 for assistance with data collection. The study is funded by the joint efforts of The Michael J. Fox Foundation for Parkinson’s Research (MJFF) and Aligning Science Across Parkinson’s (ASAP) initiative. MJFF administers the grant ASAP-000350 (to J.H.H. and M.L.) on behalf of ASAP and itself. The research was also supported by National Institute of General Medical Sciences, NIH, R01 GM111730 (J.H.H.), the National Health and Medical Research Council (NHMRC) GNT1106471 (M.L), and the Australian Research Council (ARC) Discovery Project DP200100347 (M.L). Use of the Stanford Synchrotron Radiation Lightsource, SLAC National Accelerator Laboratory, is supported by the U.S. Department of Energy, Office of Science, Office of Basic Energy Sciences under Contract No. DE-AC02-76SF00515. The SSRL Structural Molecular Biology Program is supported by the DOE Office of Biological and Environmental Research, and by the National Institutes of Health, National Institute of General Medical Sciences (P30GM133894). The contents of this publication are solely the responsibility of the authors and do not necessarily represent the official views of NIGMS or NIH.

## Competing interest statement

J.H.H. is a cofounder of Casma Therapeutics and receives research funding from Casma Therapeutics, Genentech and Hoffmann-La Roche. M.L. is a member of the Scientific Advisory Board of Automera. The other authors declare that they have no competing interests.

---

**Supplemental Figure 1.**
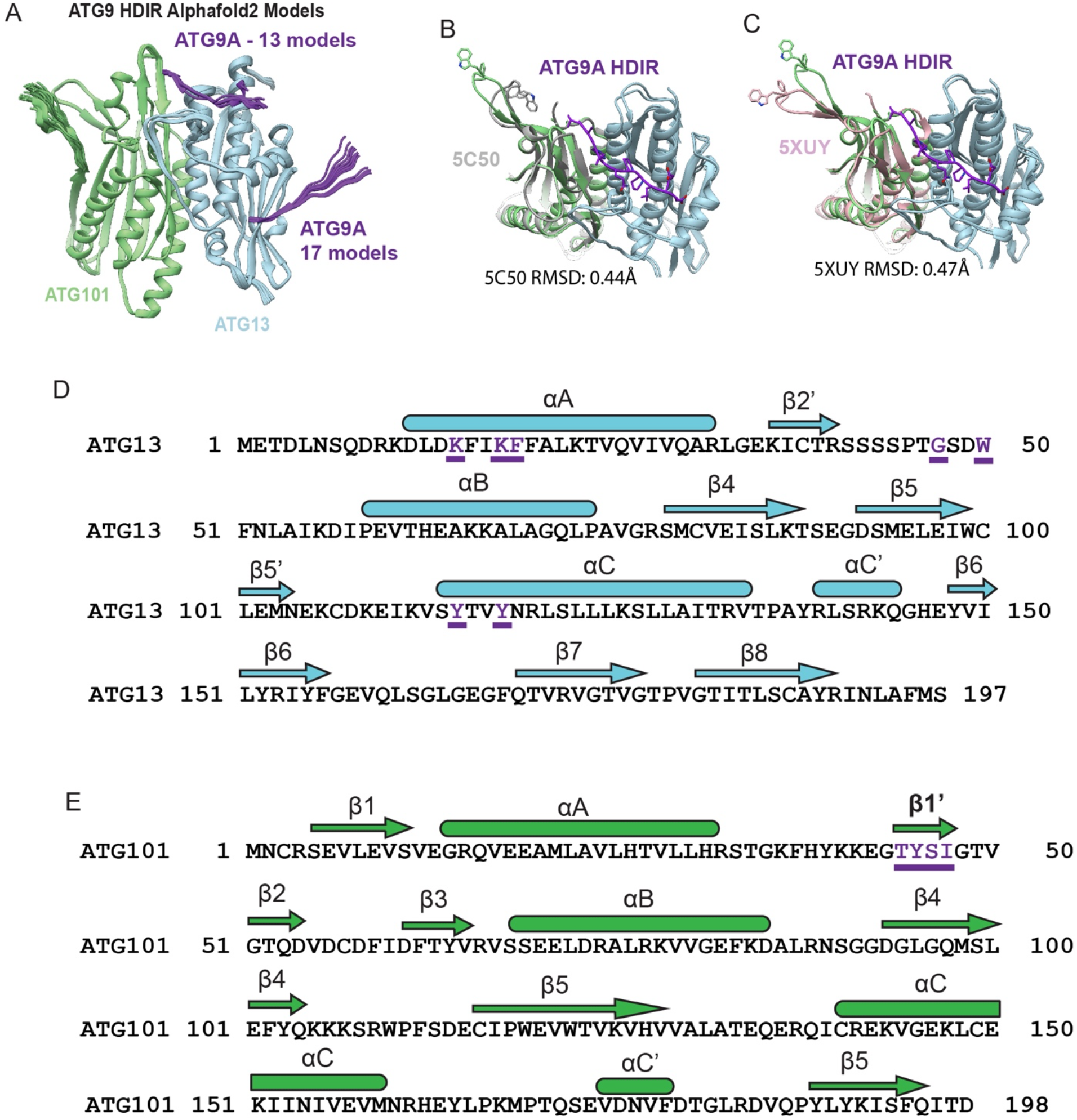
Structural data for ATG9 HDIR-ATG101:ATG13. A) Overlaid Alphafold2 models colors as in Figure 2 B) and C) Comparison of the ATG9 HDIR-ATG101:ATG13 to apo ATG101:ATG13 structures, PDB:5C50 and PDB:5XUY, respectively. D) Secondary structure of ATG13^HORMA^ with residues that interact with ATG9 HDIR shown in purple E) Secondary structure of ATG101 with residues that interact with ATG9 HDIR shown in purple

**Supplemental Figure 2.**
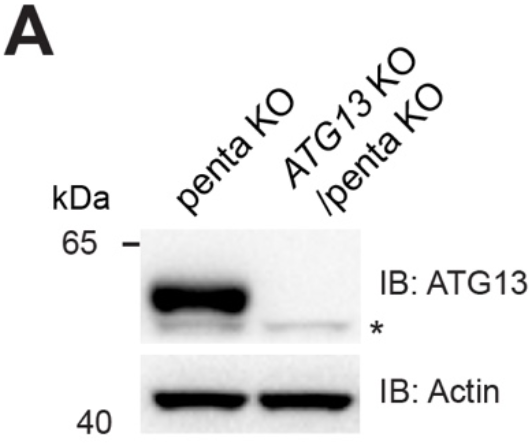
Western blot for *ATG13* KO in penta KO cell line. A) Immunoblotting of ACTIN and ATG13 in *ATG13* KO/penta KO cell line

